# Global-scale structure of the eelgrass microbiome

**DOI:** 10.1101/089797

**Authors:** Ashkaan K Fahimipour, Melissa R Kardish, Jonathan A Eisen, Jenna M Lang, Jessica L Green, John J Stachowicz

## Abstract

Plant-associated microorganisms are essential for their hosts' survival and performance. Yet, most plant microbiome studies to date have focused on terrestrial plant species sampled across relatively small spatial scales. Here we report results of a global-scale analysis of microbial communities associated with leaf and root surfaces of the marine eelgrass *Zostera marina* throughout its range in the Northern Hemisphere. By contrasting host microbiomes with those of their surrounding seawater and sediment communities, we uncovered the structure, composition and variability of microbial communities associated with *Z. marina*. We also investigated hypotheses about the mechanisms driving assembly of the eelgrass microbiome using a whole-genomic metabolic modeling approach. Our results reveal aboveground leaf communities displaying high variability and spatial turnover, that strongly mirror their adjacent coastal seawater microbiomes. In contrast, roots showed relatively low spatial turnover and were compositionally distinct from surrounding sediment communities — a result largely driven by the enrichment of predicted sulfur-oxidizing bacterial taxa on root surfaces. Metabolic modeling of enriched taxa was consistent with an assembly process whereby similarity in resource use drives taxonomic co-occurrence patterns on belowground, but not aboveground, host tissues. Our work provides evidence for a core *Z. marina* root microbiome with putative functional roles and highlights potentially disparate processes influencing microbiome assembly on different plant compartments.

## 1 Introduction

The health and performance of plants are often modulated by their associated microbiomes. Colonization of above- and belowground plant tissues by microorganisms from surrounding environments initiates interactions that are essential for plant productivity (Fürnkranz et al, 2008; Panke-Buisse et al, 2015), fitness (Lindow and Brandl, 2003; Haney et al, 2015) and disease resistance (Mendes et al, 2011; Berendsen et al, 2012; Wei et al, 2015). The drivers of plant microbiome structure and composition, and the ways in which plant hosts acquire microorganisms from surrounding microbial species pools, therefore have consequences for ecosystem dynamics, biodiversity and agricultural productivity (Philippot et al, 2009; Bakker et al, 2012; Turner et al, 2013; Berg et al, 2015). Recent studies have identified critical associations between host and environmental factors, and patterns of microbial community structure on plant compartments like leaves (e.g., Kembel et al, 2014; Laforest-Lapointe et al, 2016) and roots (e.g., Berendsen et al, 2012; Edwards et al, 2015). Yet, most plant microbiome studies to date have focused on terrestrial species (Turner et al, 2013; Grube et al, 2015; Laforest-Lapointe et al, 2016), while patterns in the structure and composition of microbial communities associated with marine plants remain poorly understood by comparison.

Seagrasses are the only flowering plants that live entirely in a marine environment. One widespread species, *Zostera marina* or eelgrass, in particular provides habitat for ecologically diverse and economically important ecosystems along coasts throughout the much of the Northern Hemisphere (Marba et al, 2007; Waycott et al, 2009; Duffy et al, 2015). The return of terrestrial seagrass ancestors to oceans is among the most severe habitat shifts accomplished by vascular plants (Hemminga and Duarte, 2000) and has prompted detailed study of the physiological adaptations associated with this shift (Pennisi, 2012; Olsen et al, 2016) including the tolerance of salinity and anoxic sediment conditions. *Z. marina* is therefore an ideal testbed for the study of microbial symbioses with plant hosts that uniquely exploit harsh environments. Given that human activities are changing nutrient conditions in habitats worldwide (Vitousek et al, 1997) and the central role of microorganisms in plant nutrition (Turner et al, 2013; Berg et al, 2015; Grube et al, 2015), there is a pressing need to answer basic empirical questions about microbial associates of plants like seagrasses that experience atypical abiotic conditions, including their geographic distributions, community assembly patterns and putative functional roles.

Much of our current knowledge of seagrass symbionts comes from targeted surveys of specific bacterial taxa using culture-dependent methods and microscopy under laboratory conditions, or from field studies at local or regional spatial scales (e.g., Newell, 1981; Kirchman et al, 1984; Donnelly and Herbert, 1998). These studies have generated hypotheses about key symbioses between seagrasses and their associated microorganisms owing to potential processes like nitrogen fixation and sulfide detoxification by bacteria (Donnelly and Herbert, 1998) and competition between microbes for host-supplied metabolites (Kirchman et al, 1984) on plant surfaces. While culture-independent techniques have been used to describe microbiome composition in seagrass-colonized marine sediments (Cifuentes et al, 2000; James et al, 2006; Cúcio et al, 2016), an extensive characterization of *in situ* seagrass leaf and root surface microbiomes across the host’s geographic range is still lacking, leaving potentially important but unculturable microorganisms overlooked and making it difficult to identify general patterns in seagrass symbiont community structure, taxonomic cooccurrence and community assembly.

Here we report results of a comprehensive analysis of microbial communities associated with leaf and root surfaces of individual *Z. marina* plants spanning their geographic range throughout the Northern Hemisphere. To determine the relative importances of potential microbial colonization sources, we characterized surrounding environments by sampling seawater and sediment communities adjacent to each collected seagrass host. We aimed to define the global structure, composition and variability of symbiont communities associated with *Z. marina*; contrast these communities with those of their surrounding environments; and investigate the mechanisms driving assembly of the seagrass microbiome using a whole-genomic metabolic modeling approach (Borenstein et al, 2008).

## 2 Methods

We sampled microbial communities present on the leaf and root surfaces of 129 eelgrass individuals, together with those from the surrounding seawater and sediment habitats, using the Illumina MiSeq platform to sequence amplified fragments of the V4 region of the 16S rRNA gene. This approach primarily targets environmental bacteria, but some archaeal sequences were also detected. Microbial samples were collected by the Zostera Experimental Network, *ZEN* — a global-scale collaboration between seagrass researchers (e.g., Duffy et al, 2015; http://zenscience.org/). Three leaf, root, water and sediment samples were collected from plots at each of 50 seagrass beds (Fig. 1a) using identical sampling protocols. Samples were placed into 2mL collection vials and covered in ZYMO Xpedition buffer. Root and leaf samples were acquired by collecting ten root hairs and a 2cm section of healthy green outer leaf blade respectively. Seawater samples were collected just above each plant by filtering approximately 300mL of seawater through a 0.22 micron filter and retaining filters. Finally, 0.25g of sediment was taken from 1cm under the surface using a syringe.

**Fig. 1.**
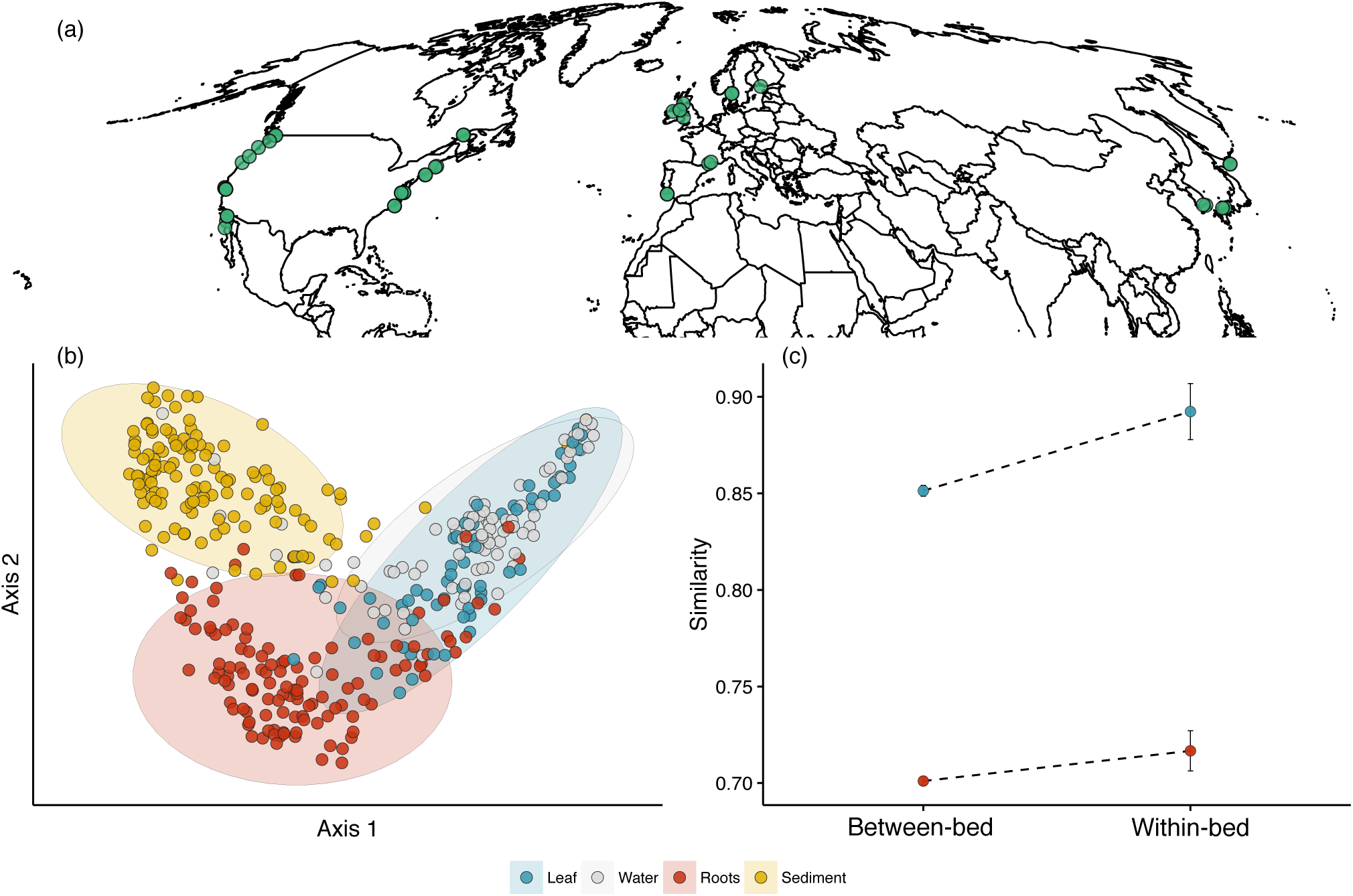
(a) Map of sampled seagrass beds. Green points represent ZEN site coordinates. (b) Ordination plots show results of a 2-dimensional PCoA of Canberra distances. Colored points correspond to sample types; blue are leaf, silver are seawater, red are root and gold are sediment samples. Ellipses represent group-specific 95% confidence intervals assuming a multivariate t-distribution. (c) Comparisons of host-environment compositional similarities within-versus between seagrass beds. Points represent mean similarities between leaves and water (blue points), and roots and sediment (red points) ± SEM.

Samples were extracted using a modified version of the MoBio PowerSoil DNA Extraction Kit Experienced User Protocol. Modifications were to remove precipitate formed by the Zymo lysis buffer and C1 solution. Tubes were incubated at 65°C for five minutes to remove precipitate and then homogenized in a bead-beater. Instead of eluting DNA in solution C6, we added 50pL of sterile, nuclease-free water to the membrane. DNA was stored at -20°C and amplified in a PCR enrichment of the V4 region of the 16S rRNA gene following a modified version of the Earth Microbiome Project’s (Gilbert et al, 2014) PCR protocol. We used the bacterial and archaeal primers 515F and 806R with an inhouse dual barcode system (see Caporaso et al, 2012). PNA blockers were used to reduce chloroplast and mitochondrial sequence products and used 1-5*μ*pL of template DNA. PCRs were cleaned with the Axygen AxyPrep Mag PCR Clean-Up Kits, quantified using Qubit and pooled with equal amounts of amplicons. Libraries were sequenced on an Illumina MiSeq generating 250bp paired end reads.

Raw sequence data were processed with QIIME (Caporaso et al, 2010) 1.9 and clustered into operational taxonomic units (OTUs) at > 97% similarity using the UCLUST algorithm (Edgar, 2010) against the *Green-Genes* version 13.8 reference database. To ensure adequate sampling depth, we omitted several samples from our analyses because they contained fewer than 1000 sequences after quality control, retaining data from 123 plants in total. We also excluded all 16S sequences identified as chloroplasts or mitochondria. The resulting OTU counts were normalized using the trimmed mean of M values (TMM) method (Robinson and Oshlack, 2010), which was chosen due to its improved sensitivity for detecting differentially abundant taxa (see below) compared to rarefaction (McMurdie and Holmes, 2014).

### Statistical analyses

Microbial community compositional and phylogenetic dissimilarities (i.e., *β*-diversities) between host and environmental samples were calculated using the Canberra and normalized unweighted UniFrac (Lozupone and Knight, 2005) distance measures respectively. Canberra distances were calculated for Hellinger-transformed normalized abundances, whereas the UniFrac measure quantifies phylogenetic distinctness of different communities based on phylogenetic relationships between OTUs that are present. Dissimilarities of host and environmental samples were visualized using unconstrained principal coordinate analysis (PCoA). Effect sizes of dissimilarities between seagrass microbial and environmental communities were quantified using a permutational analysis of similarities (ANOSIM), and differences in group variances were tested using a multivariate homogeneity of groups dispersions analysis (betadisper; Anderson, 2006) with pairwise comparisons made with ANOVA and Tukey’s Honest Significant Differences test. *β*-diversity analyses were conducted using the *vegan* package in the statistical programming environment R (R Core Team, 2016).

We compared community compositions of host and environmental samples at the scale of the seagrass bed, to test the hypothesis that host-associated microbiomes were more similar to their adjacent environmental communities (i.e., within-bed comparison) than to others (i.e., between-bed comparison). We did this using a Monte Carlo bootstrapping approach, similar to Song et al (2013), following ordination analyses. To accomplish this we first computed the distances between group centroids of host samples taken from the same seagrass bed and the centroids of their corresponding environmental samples. We then determined whether host-associated microbial communities were more similar to their adjacent environment than to others by comparing inter-centroid distances against the distributions generated from 1000 permutations of the randomized dataset. Performing *β*-diversity analyses for both the Canberra and UniFrac distance measures allowed us to determine the degree to which microbiomes found on different compartments of the same host differed from one another and those of their surrounding environments, both compositionally and phylogenetically.

Environmental sources of microorganisms detected on seagrass leaves and roots were estimated by training a Bayesian source tracking classifier (*Source-Tracker*; Knights et al, 2011) on the set of water and sediment microbiome samples from each sampled coastline before testing the model on corresponding host samples. The model assumes that host communities comprise a combination of colonists that originated from known and unknown exogenous sources and, using a Bayesian approach, estimates the fraction of OTUs detected on each leaf and root surface that originated from water, sediment or unknown habitats. We used the estimates from this classifier to perform a guided differential abundance analysis for the two host compartments to identify OTUs that were significantly enriched or depleted on leaves and roots relative to their primary putative colonization source. We did this by fitting generalized linear models with negative binomial error distributions to TMM-normalized OTU counts and identifying differentially abundant taxa on host samples using a likelihood ratio test. We focused subsequent analyses on OTUs that were significantly host-enriched (Benjamini-Hochberg adjusted *P* < 0.01), as these taxa represent portions of the microbiome that were most likely to be actively selected for by the host (Burns et al, 2015).

Potential drivers of the acquisition of enriched taxa were investigated using metabolic modeling (Borenstein et al, 2008; Levy and Borenstein, 2013) of these taxa or their closest relatives with fully-sequenced genomes in the NCBI reference database (Pruitt et al, 2005). Namely, we sought to determine whether enriched taxa that are predicted to utilize similar metabolite resources on eelgrass surfaces co-occurred more or less frequently than expected by chance. To accomplish this, we conducted a BLAST sequence similarity search (Altschul et al, 1990) comparing each enriched OTU to a database of 16S sequences for prokaryotic taxa with whole genome sequences in NCBI, compiled by Mendes-Soares et al. (2016). The ModelSEED framework (Devoid et al, 2013) was used to reconstruct and gap-fill models for the genomes most similar to eelgrass-enriched OTUs. Metabolic models were represented as topological networks where nodes denote chemical compounds and directed edges connect chemical reactants to products. Using these networks, each OTU’s seed set (Borenstein et al, 2008) — the minimal set of compounds an organism exogenously acquires to synthesize all others in its metabolic network — was calculated as a proxy for its nutritional profile (Levy and Borenstein, 2013) using a previously published graph-theoretic method (Borenstein et al, 2008).

After computing each enriched OTU’s seed set, a competitive dissimilarity matrix **C** was generated, which contained elements *C_ij_* representing the pairwise uniqueness of taxonomic resource profiles, defined as the fraction of seeds in the seed set of OTU *i* not shared with j. Values of 1 in this matrix indicate no overlap between the seed sets of two OTUs (i.e., no predicted resource overlap) whereas a value of 0 indicates that two OTUs had identical seed sets. The relationship between co-occurrence dissimilarity (measured as Jaccard distances) and OTU competitive dissimilarity (*C_ij_* values) matrices were assessed for leaf- and root-enriched taxa using Mantel tests with 1000 matrix permutations. If enriched taxa utilize similar predicted resources, then we would expect a positive correlation between the Jaccard and **C** distance matrices. Such patterns are consistent with a habitat filtering community assembly mechanism whereby organisms that require a set of resources tend to co-occur in environments with those resources (Levy and Borenstein, 2013).

## 3 Results

We identified 23,285 microbial operational taxonomic units (OTUs, sequences binned at a 97% similarity cutoff) on eelgrass host surfaces, an average of 492.3 ± 40.3 OTUs (mean ± SEM) per leaf sample and 1304.6 ± 62.8 per root sample. A higher number of OTUs were detected in environmental samples on average, observing a mean of 589.9 ± 64.2 OTUs in seawater samples and 1767.4 ± 66.3 in sediment samples. A larger proportion of the taxa detected on leaves were rare compared to those on roots — 92.5% of the OTUs detected on leaves were observed on fewer than five leaves compared to 75% for roots — consistent with the occurrence of higher taxonomic turnover on aboveground plant compartments. Indeed, *β*-diversity analysis of seagrass symbiont communities revealed major differences in above-versus belowground seagrass microbiomes and their relationships with the surrounding environment (Fig. 1). Taxonomic composition of the seagrass leaf microbiome was quite variable and strongly resembled that of seawater, whereas root communities were relatively similar to one another and fairly distinct from sediment (Fig. 1a). The *Z*. *marina* leaf microbiome was more similar to that of seawater (ANOSIM of Canberra distances; *r* = 0.15, adjusted *P* < 0.001) than the root microbiome was to sediment communities (ANOSIM; *r* = 0.56, *P* < 0.001; compare *r* statistics).

The taxonomic composition of leaves and seawater microbiomes were more similar within seagrass beds than between them (Fig. 1b, blue points; *P* = 0.017), a result that is consistent with a seagrass leaf driven by the microbial composition of the local ocean environment. In contrast, we did not detect a higher degree of compositional similarity between roots and sediment sampled from the same seagrass bed relative to other beds (Fig. 1b, red points; *P* = 0.24), suggesting more homogenous microbiome taxonomic compositions at the global scale. These results were recapitulated by a multivariate dispersion analysis, which revealed aboveground host and environmental community compositions that exhibited variances that were indistinguishable from one another (betadisper pairwise leaf and water comparison; *P* = 0.96), and root microbiomes that were globally less variable compared to sediment communities (betadisper; *P* < 0.001). Analyses of unweighted UniFrac distances revealed similar qualitative results for patterns in phylogenetic *β*-diversity *(Supplementary Information*).

### Environmental sources of seagrass-associated microorganisms

We estimated the relative contributions of sediment, seawater and unknown environmental sources for individual samples of seagrass leaf and root microbiomes using the Bayesian *SourceTracker* classifier (Knights et al, 2011). The model estimates that seawater is the primary source of colonists for seagrass leaves (Fig 2a; median proportion of water-sourced OTUs = 0.8), with many leaf samples appearing nearly entirely water-sourced (Fig. 2a, dark blue shaded area). Roots were estimated to be primarily sourced from sediment (Fig. 2b; median proportion of sediment-sourced OTUs = 0.51). Although the communities on some roots were predicted to originate nearly completely from sediments, most appeared to receive colonists from both above- and belowground environments (Fig. 2b, dark red area).

**Fig. 2.**
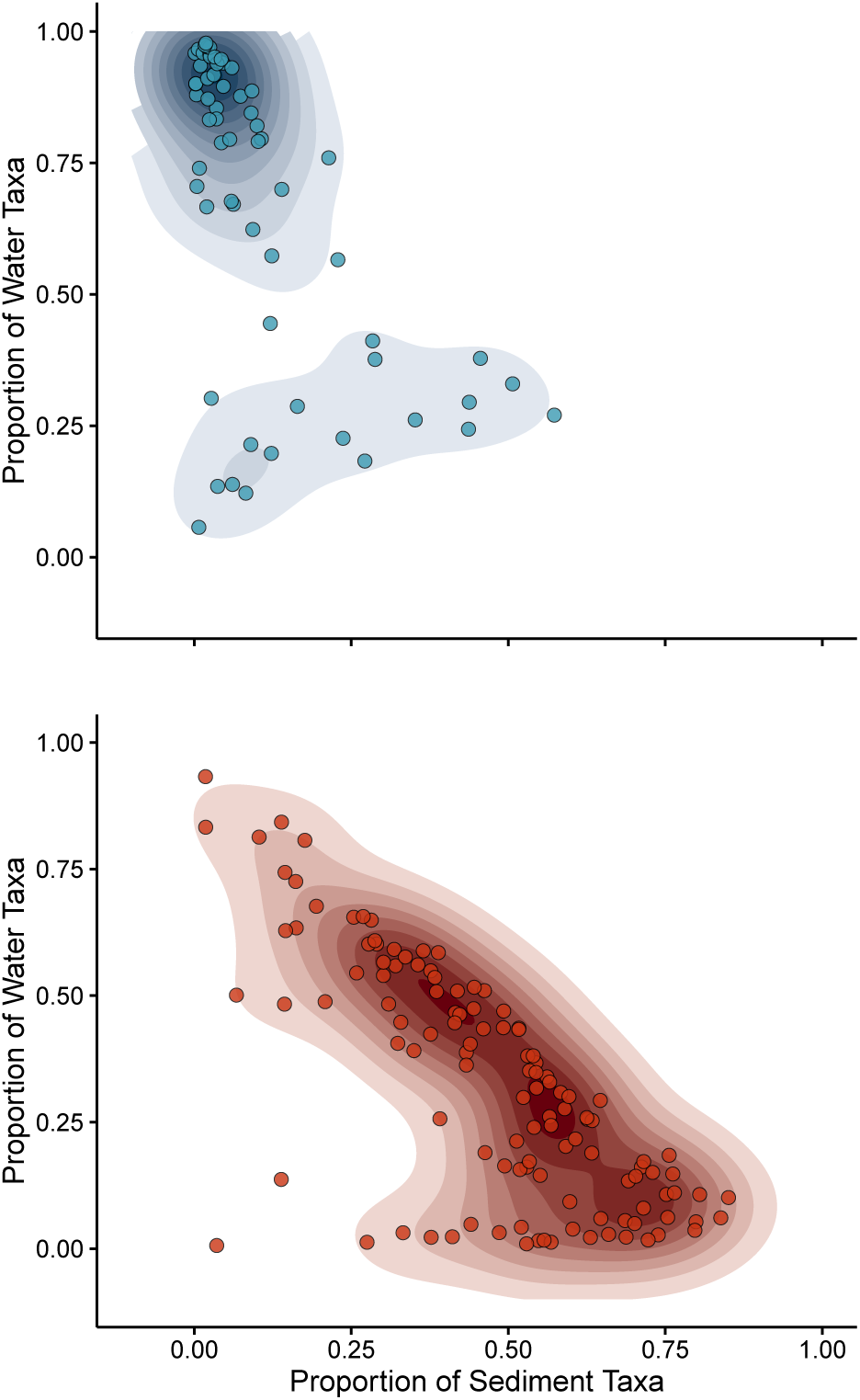
Results of *SourceTracker* analysis for (a) leaf and (b) root samples, where points represent individual microbial communities. Colors are the same as in Fig. 1. Contours are shaded according to a 2d Gaussian kernel used for density estimation, where darker shades represent denser clusters of data points.

We used the estimates from source tracking to perform a guided differential abundance analysis for each of the two plant compartments (i.e., leaves and roots), to identify OTUs that were significantly enriched or depleted on hosts relative to their abundances in the primary putative colonization source. We observed 39 enriched and 126 significantly depleted OTUs on *Z. marina* leaves relative to water (Fig. 3a), revealing an aboveground host compartment in which fewer than 10% of detected taxa exhibited patterns in normalized abundance that differed from those observed for seawater communities. Leaf-enriched taxa were largely represented by members of the *Gammaproteobacteria, Planctomycetia, Flavobacteriia* and *Betaproteobacteria* classes (Fig. 4, blue columns). In contrast, we detected 510 enriched and 1,005 depleted OTUs on sea-grass roots (Fig. 3b), consistent with a higher degree of host recruitment and higher selectivity against particular environmental microorganisms on belowground seagrass tissues; 25% of taxa detected on roots exhibited patterns in normalized abundance that differed from those observed in sediments. Notably, 50 of these root-enriched OTUs (c.a. 10%) clustered onto the genus *Sulfurimonas*, of which most of the cultured isolates are sulfide-oxidizers (Han and Perner, 2015). Moreover, 25% of root-enriched OTUs were members of the *Desulfobulbaceae, Desulfovibrionaceae, Desulfuromonadaceae* or *Desulfobacteraceae* families or the *Arcobacter* genus, highlighting the acquisition of a diverse set of OTUs related to taxa involved in sulfur metabolism by belowground tissues as a potentially key process for marine angiosperms.

**Fig. 3.**
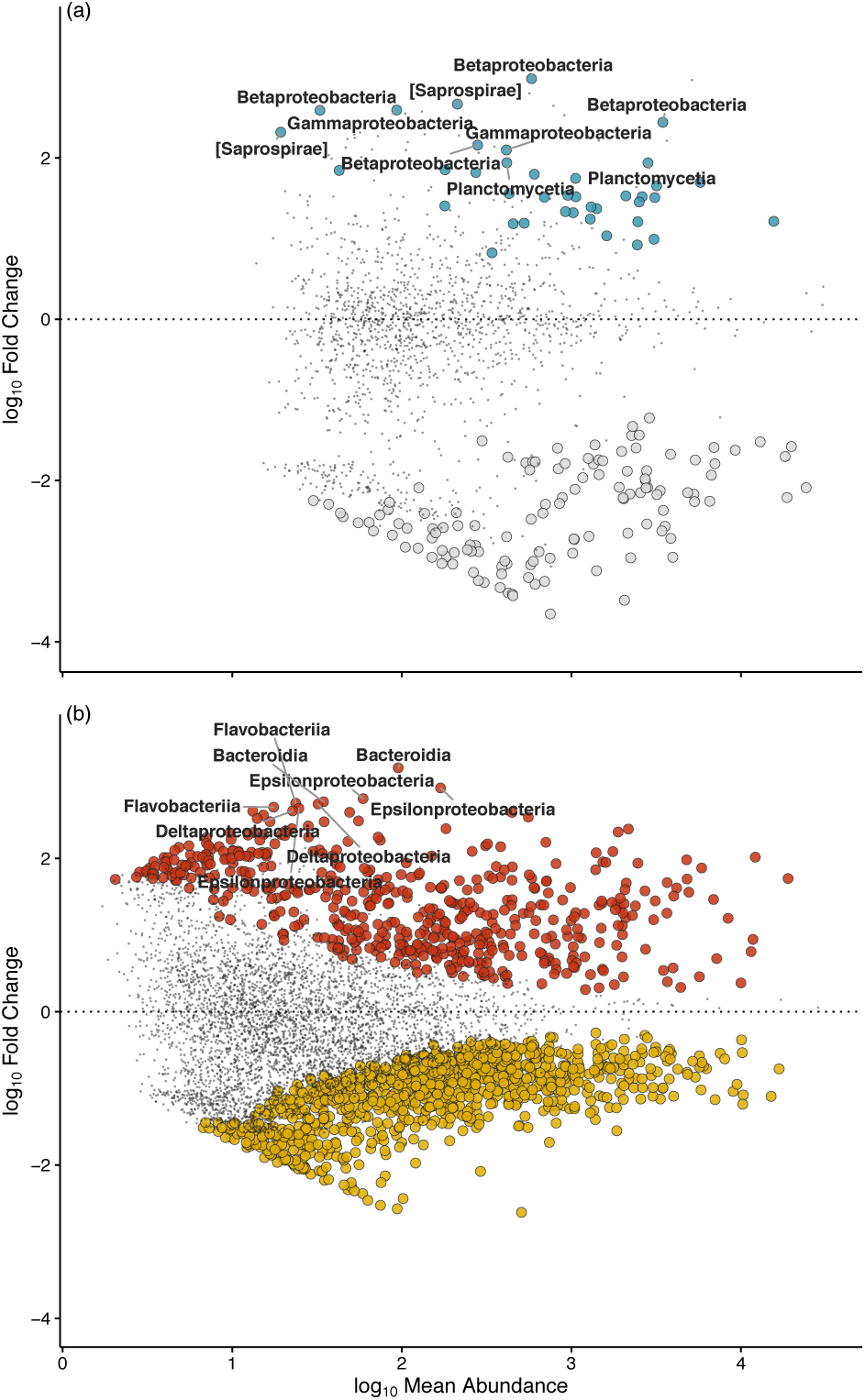
Host compartments are enriched and depleted for certain OTUs. (a) Enrichment and depletion of OTUs detected on leaves compared to the seawater environment as determined by differential abundance analysis. Each point represents an individual OTU, and the position along the y axis represents the abundance fold change relative to the primary source environment. Colors are the same as in Fig. 1; significantly enriched and depleted OTUs are colored blue and silver respectively. (b) Results of differential abundance analysis for OTUs detected on roots compared to the sediment environment. Significantly enriched and depleted OTUs are colored red and gold respectively. The taxonomic class of the top ten most enriched taxa are labelled for reference.

**Fig. 4.**
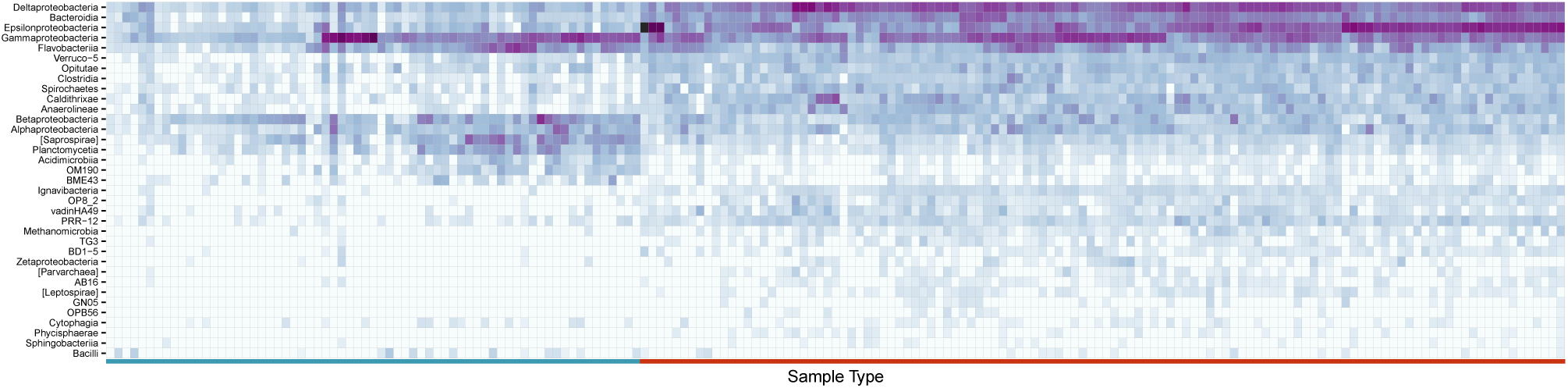
Heatmap showing the taxonomic compositions of enriched taxa, aggregated at the class-level, on leaves and roots. Darker shades of purple correspond to higher mean abundances of OTUs in each class. Leaf and root samples are differentiated on the x-axis by blue and red markers respectively. White tiles indicate those taxa were not detected in particular samples. Matrix seriation was accomplished using a hierarchical clustering algorithm with an average linkage method.

### Metabolic models of host-enriched taxa support hypotheses about seagrass microbiome assembly

We sought to investigate potential mechanisms underlying the enrichment of taxa on seagrass surfaces, by investigating whether host-enriched taxa that are predicted to utilize similar metabolite resources (i.e., higher predicted strength of competition) on seagrass surfaces cooccur more or less frequently than expected by chance through metabolic modeling of leaf- and root-enriched taxa. Leaf- and root-enriched taxa exhibited median similarities to the 16S sequences of their most similar genomes of 91.6% and 92.9% respectively. We did not detect a significant relationship between dissimilarity in predicted resource use and OTU co-occurrence for enriched taxa on leaves (Fig. 5a; Mantel *P* = 0.36). However, a significant positive relationship was observed among root-enriched OTUs (Fig. 5b; Mantel *P* = 0.006), indicating that taxa with higher resource overlap co-occur more frequently, on average. Importantly, this relationship held when we accounted for pairwise phylogenetic branch lengths between OTUs (partial Mantel *P* = 0.002), indicating that metabolic modeling was not simply recapitulating phylogenetic relationships between taxa (Levy and Borenstein, 2013).

**Fig. 5.**
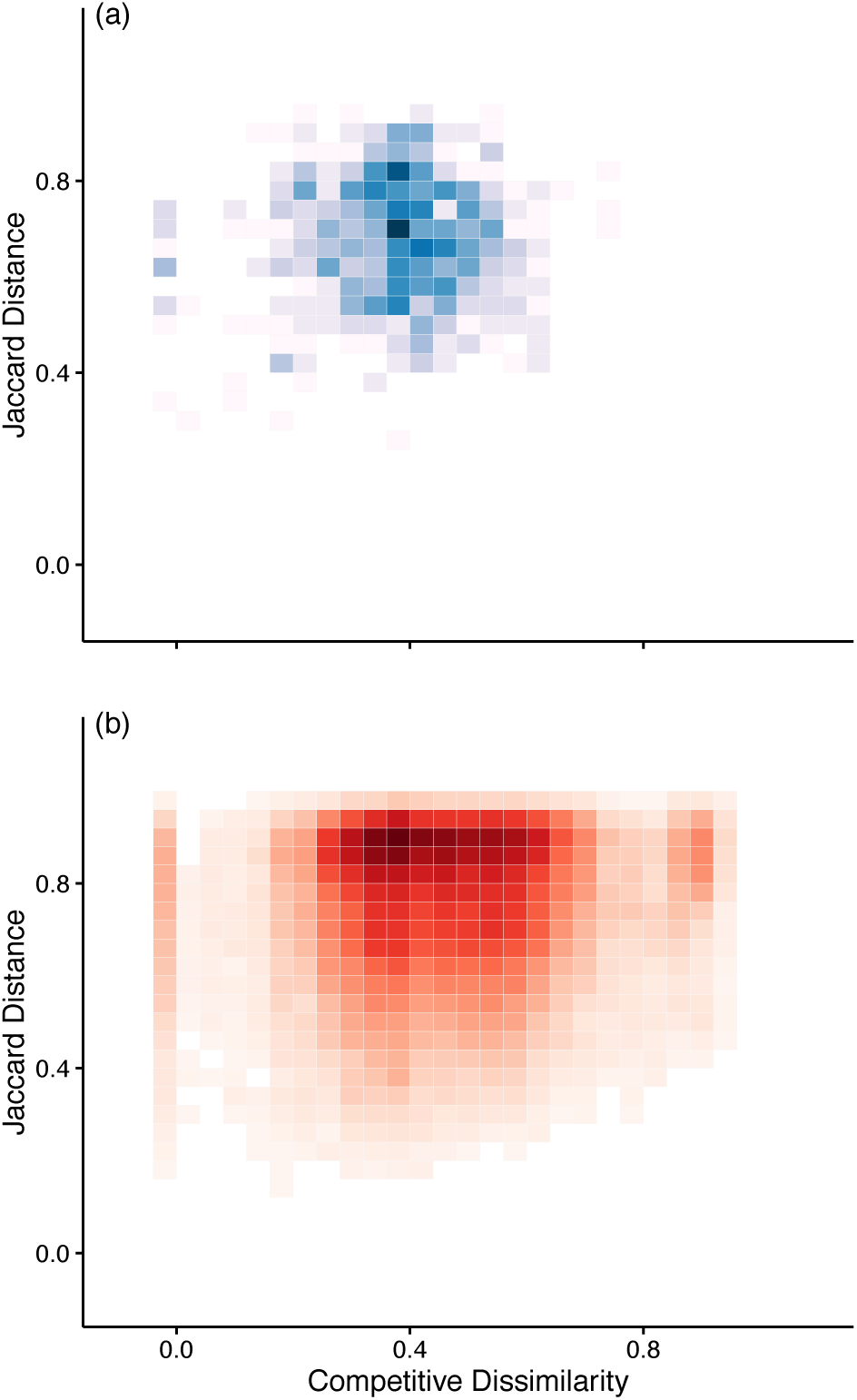
Relationships between OTU co-occurrence (Jaccard distance) and competitive dissimilarity matrices for host-enriched taxa. Matrix comparisons were visualized using 2d histograms which show the distributions of values in a data set across the range of two quantitative variables, where darker colors represent higher frequency bins. (a) Relationship between binned leaf-enriched OTU Jaccard distance and competitive dissimilarity matrices. (b) Relationship between root-enriched OTU Jaccard distance and competitive dissimilarity matrices. A positive relationship on roots (P = 0.006) is consistent with a habitat filtering community assembly mechanism.

## 4 Discussion

Our global study of the Zostera marina eelgrass microbiome revealed a high degree of similarity between leaf and seawater communities compared to root surfaces, whose taxonomic and phylogenetic compositions were less heterogenous than, and more distinct from, the surrounding sediment (Fig. 1). As very few studies describe the structure of microbiomes associated with aquatic plant surfaces compared to terrestrial species (Crump and Koch, 2008), observations of terrestrial plants serve as an important reference. Our results identify notable contrasts in the structure of the eelgrass microbiome compared to those observed on well-studied terrestrial species. For instance, the commonality between seagrass leaf and adjacent seawater microbiome compositions differs from relationships observed for terrestrial plant leaves, which appear distinct from the microbial communities observed from air sampling (Bowers et al, 2009; Redford et al, 2010; Vorholt, 2012; Womack et al, 2015). Eelgrass leaves in our study exhibited microbiome compositions that strongly mirrored their surrounding seawater communities (Fig. 1b). Notably, *Z. marina* has lost genes for the production of volatile terpenes and lack stomata on leaves (Olsen et al, 2016), raising the possibility that seagrass leaves lack many of the characteristics of terrestrial plants (e.g., localized gas exchange via stomata, chemical defense and communication) thought to influence the structure of their associated leaf microbiomes.

The widespread success of seagrasses has occurred despite environmental challenges. In particular, organic matter accumulation within coastal sediments causes toxic sediment sulfide conditions for vascular plants (Jørgensen, 1982; van der Heide et al, 2012). The most abundant of the root-enriched microbial taxa detected in the present study clustered onto the genus *Sulfurimonas*, which accounted for approximately 10% of all root-enriched OTUs. All but one of the previously isolated strains of *Sulfurimonas* can oxidize sulfide and produce sulfate as an end product, suggesting that the recruitment of these bacteria may be critical for host tolerance of coastal marine habitats. Oxidation of sulfide and its precipitation as nontoxic S^0^ on the inner wall of the host’s aerenchyma tissue has previously been attributed to host detoxification mechanisms like the leakage of oxygen from root tips (Hasler-Sheetal and Holmer, 2015). The enrichment of *Epsilonproteobacteria* like *Sulfurimonas* on root surfaces, and the consistency of this pattern at the global scale, however adds further support to the hypothesis that microbial symbioses with particular taxa facilitate seagrass hosts’ management of sulfide toxicity in coastal beds. Indeed, abundant bacteria that are predicted sulfur-oxidizers have been observed in marine sediments attached to seagrass roots (Cucio et al, 2016), and T-RFLP community profiling (Liu et al, 1997) of root surfaces in a single European seagrass bed has suggested similar patterns in *Epsilonproteobacteria* community dominance (Jensen et al, 2007). Results of our metabolic modeling suggests that hosts may enrich for these microorganisms in part via the supply of particular metabolic compounds to belowground plant compartments. Although predictions from metabolic modeling are consistent with prior studies of host-supplied metabolites on seagrass surfaces (e.g., Kirchman et al, 1984), these predictions do have limitations and should be interpreted as hypotheses. The metabolic models analyzed herein are derived from 16S sequences and involve automated metabolic network reconstruction (Devoid et al, 2013; Mendes-Soares et al, 2016). This approach may be less accurate than manual curation of metabolic models (Mendes-Soares et al, 2016), but automation permits the analysis of a large number of microbial taxa that would otherwise be intractable and indeed reflects a large proportion of the metabolic capabilities of these organisms.

Prior research has documented a positive relationship between seagrass biomass production rates and the density of sulfide-consuming *Lucinid* clams in seagrass beds, owing to the hypothesized *in situ* reduction of sulfide concentrations by symbiotic bacteria housed in clam gills (van der Heide et al, 2012). However, in a meta-analysis of temperate seagrass beds only 50% of sampled beds contained *Lucinid* bivalves, and clam density was low in these beds relative to tropical sites (van der Heide et al, 2012). Thus, temperate seagrasses must either be more tolerant of sulfides or have alternative means of detoxification. Physiological host processes like oxygen leakage from roots (Hasler-Sheetal and Holmer, 2015) certainly contribute to sulfide oxidation, but our data suggest a role for microorganisms directly associated with eelgrass; *Sulfurimonas* bacteria occurred in all but one root sample. Experimental efforts are therefore needed to quantify the magnitudes of sulfur metabolism from these disparate processes (oxygen leakage, lucinid bivalves, root associated bacteria) under different biotic and abiotic conditions, in order to uncover the relative importance of host-versus mutualism-based strategies for tolerating toxic sulfide concentrations by vascular plants in marine sediments.

Seagrasses and their ecosystems have been the subject of a great amount of research covering many topics including ecology and biogeography (Duffy, 2006), evolution (Chen et al, 2012), physiology (Pennisi, 2012) and genetics (Olsen et al, 2016). Here, we have provided a global-scale characterization of the microbial communities associated with *Z. marina* seagrasses by contrasting host samples with those of their surrounding environments across the entire Northern Hemisphere. We hope that this will encourage researchers to study the microbiomes of other plant hosts across their geographic ranges, as such broad scale studies produce the empirical knowledge needed to develop a deeper understanding of microbial roles in the ecology and evolution of plants and the ecosystems that depend on them.

## Acknowledgements

We thank the ZEN partners for their assistance with data collection. We also thank Steven Kembel, James Meadow and Roxana Hickey for helpful discussions. This study was supported by a grant from the Gordon & Betty Moore Foundation to JAE, JML, JLG and JJS.

